# Early Changes in Respiration and Motor Activity after Spinal Cord Injury Predict Pain-Related Outcomes in Mice

**DOI:** 10.1101/2025.08.29.672856

**Authors:** Austin Chuang, Shawn Hochman, Donald J. Noble

## Abstract

Spinal cord injuries (SCIs) may lead to the emergence of chronic pain many weeks after injury. Using the thoracic contusion model of SCI-induced neuropathic pain, we investigated whether early changes in mouse respiration or motor activity could predict and differentiate emergent dysfunction with focus on pain. We measured respiratory rate (RR) and movement (motor activity) in freely behaving mice before and at several time points following SCI. We then assessed behavioral signs of pain or thermal dysregulation by testing evoked hindpaw withdrawal responses to mechanical and thermal heating stimuli and temperature preferences at four weeks after injury. For 2-3 days after injury, mice exhibited sharp decreases in movement and RR variability, but these two parameters were uncorrelated within animals. Mice showed signs of mechanical and thermal hypersensitivity and preferences for warmer temperatures four weeks after injury. Interestingly, mice that moved the least 1 day after SCI preferentially underwent hindpaw mechano-sensitivity (*r* = 0.67, *p* =.036), whereas mice with large decreases in RR variability that recovered by 8 days post-injury preferred higher temperatures in the thermal preference test (*r* = 0.79, *p* =.007). Thus, early changes in movement and RR variability may differentially predict future hypersensitivity and thermal dysfunction. More broadly, early post-injury physio-behavioral events could inform novel interventions to mitigate subsequent emergent dysfunction after SCI.

## Introduction

A spinal cord injury (SCI) is a serious neurological disease that is difficult to treat and may cause lifelong impairments. An estimated 80% of SCI patients suffer from uncurable neuropathic pain due to incompletely understood biological mechanisms and generally inadequate pharmacological treatments.^1-3^ Neuropathic pain develops from damage to the somatosensory nervous system.^4^ Regions that maintain partial sensory functionality often lead to allodynia, a type of hypersensitivity characterized by a painful response to typically non-painful stimuli such as a light touch or mild temperature.^5^ More broadly, neural injury-induced pain frequently co-occurs with increases in sympathetic activity.^6-10^ Nociceptive stimuli may increase heart rate (HR) and respiratory rate (RR) by activating the sympathetic nervous system.^11-14^ High RR predicts numerous negative cardiopulmonary outcomes,^15,16^ and may have greater predictive potential for early detection of severity and risks of a given injury.^17^ Recently, studies have observed that higher resting RR and RR variability (RRVAR) after lower thoracic SCI are correlated with physiological and behavioral measures of pain in rodents.^18,19^ Whether interventions involving normalization of respiration ameliorate pain outcomes following SCI remains to be determined.^20^

In addition to SCI patients experiencing pain-related symptoms, changes in motor function are also highly prevalent,^21^ and may be related to the emergence of allodynia. Although our understanding of the mechanisms causing neuropathic pain after SCI is limited,^3,22^ the amount of spared tissue post-SCI may correspond to increased spinothalamic function and neuropathic pain.^23^ Magnitude of spared mid-sagittal tissue bridges also correlated with the extent of motor recovery in SCI patients.^24^ There is a positive correlation between the severity of neuropathic pain and overall functional motor recovery in SCI patients.^25^ Not yet studied is whether an earlier motor recovery may predict subsequent emergence of neuropathic pain.

We hypothesized that SCI leads to early differentiable changes in function that are predictive of subsequent dysfunction. Studies were undertaken using a lower thoracic cord contusion SCI model of neuropathic pain^26-28^ and motor impairment.^29,30^ We quantified early changes in respiratory and motor activity as possible predictors of SCI pain-related outcomes. Neuropathic pain was assessed through tests of reflexive mechanical (von Frey) and thermal (Hargreaves) hypersensitivity, with additional testing for thermal dysregulation (Thermal Place Preference).

## Materials and Methods

All protocols conformed to the Guidelines for the Care and Use of Laboratory Animals of the National Institutes of Health and were approved by the Emory University Institutional Animal Care and Use Committee.

### Subjects and Surgical Procedures

Adult male and female C57BL/6J mice (The Jackson Laboratory, #000664) were used. Mice were anesthetized with isoflurane (5%, gas; lowered to 2-3% upon reaching stable anesthesia). Under sterile conditions, the spinal cord was exposed following skin incision and dorsal laminectomy, and the injury was administered as previously described.^31^ Briefly, mice received a moderate (∼70kDyne) impact contusion SCI (IH-0400 Impactor, Precision Systems and Instrumentation, Fairfax Station, VA, USA) delivered to the lower thoracic (T) spinal cord at level T10. The overlying skin was sutured shut and the wound area treated with a topical ointment. Mice were weighed daily following surgery and their bladders expressed by researchers twice daily for the duration of experiments. Three cohorts of mice were used in this study. The first cohort (n=7) was sacrificed nine days post-surgery for use in other experiments. As a result, tests occurring at the four-week post-surgical time point only involved the second (n=6) and third (n=4) cohorts. All mice were administered meloxicam [5 mg/kg, SC] and buprenorphine [0.05 mg/kg, SC] prior to surgery for acute pain management, then left to recover on a heated pad. The same meloxicam dosage was delivered twice daily for 2 days following surgery. Sterile saline [0.9%, IP], was administered daily for the first 48 h after surgery with subsequent injections given as needed. Findings presented at 1-2 dpo (meloxicam administered) and 3 dpo (no meloxicam) were consistent, suggesting that analgesic effects did not play a major role.

### Breathing and Motor Activity

Mice were placed in square enclosures with two electric field (EF) sensors (EPIC, Plessey Semiconductors, Plymouth, UK; now commercially available at Level 42 AI, Inc, Mountain View, CA) affixed to the side of each enclosure to non-invasively record RR and its variability in mice acutely after injury (**Figure 1A**).^32,33^ Sensor recordings were captured as voltage traces through a customized interface in LabVIEW (National Instruments, Austin, TX). A lowpass Chebyshev filter at 12Hz was used to remove electrical noise. Resting RR, RRVAR, and movement were quantified via threshold-based event detection in Clampfit (bandpass filter 1-7Hz; Molecular Devices, San Jose, CA) in combination with derived spectrograms (**Figure 1B**). Before surgery, each mouse underwent a 1.5-hour acclimation session in the same enclosures as above, with the final 30-minute period recorded used to calculate the baseline RR, RRVAR (the standard deviation of RR),^18^ and time spent moving. This design was chosen following previous testing to ensure reliability of environmental acclimation. The day after their baseline recordings were taken, mice received the SCI, with subsequent post-acclimation 30-minute recording sessions beginning at 1 day post-SCI. Post-surgical time-points included the first three days post injury (acute time points), day 8 post injury (subacute time point), and four weeks after injury (chronic timepoint). We were unable to obtain recordings from n=4 mice at 2 days post operation (dpo) due to data capture error resulting in n=13 SCI mice at this time point (n=17 for others).

**Figure 1:**
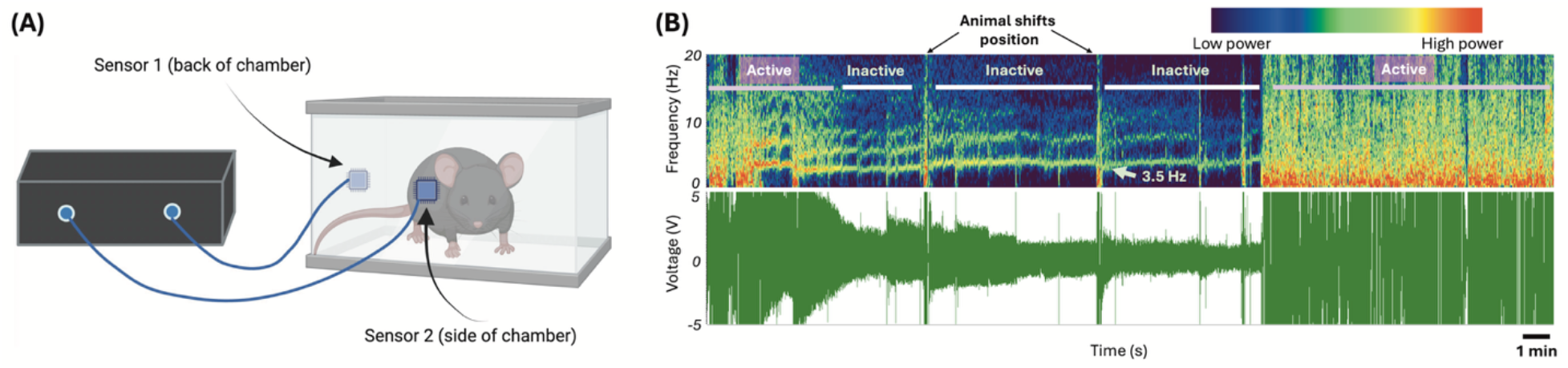
Electric field sensor recording apparatus with example captured data. **(A)** Mice were placed in acrylic chambers allowing for full range of movement. Each mouse chamber had two affixed sensors that translate activity-based changes in the electric field to changes in voltage. **(B)** Example raw voltage recording (*bottom*) and transformed frequency spectrograms generated in Spike2 (Cambridge Electronic Design, Milton, England, *top*). Note the discrete power bands associated with respiration rate (frequency) when mouse is at rest (inactive), and that active motor epochs have larger voltage responses with broad increases in power across spectral frequencies displayed. Time spent moving, a spontaneous motor behavior, was calculated from spectrograms as total trial time minus total length of resting respiration epochs. Results were verified using simultaneous video capture in a subset of mice.

### Mechanical and Thermal Sensitivity Tests

The von Frey hindpaw mechanical sensitivity test was undertaken at baseline (before surgery) and at the 4-week post-SCI time point, at which many studies have shown full development of mechanical hypersensitivity associated with neuropathic pain.^19,31,34^ For 3 days, mice were habituated to the behavioral testing room, including the red acrylic chambers where testing was performed. On testing days, following a 30-minute acclimation period, individual animals were assayed for mechanical sensitivity according to the established up-down method,^35^ using calibrated von Frey hairs (NC12775-99, North Coast Medical, Inc., Morgan Hill, CA, USA). Right and left hindpaw withdrawal thresholds were averaged to determine overall mechanical sensitivity.

One day after the von Frey test and using the same acrylic chambers, the Hargreaves test was conducted to quantify thermal nociception.^36^ The test was run at 4-weeks post-SCI, but not at pre-injury baseline to avoid habituation effects observed in previous cohorts. A group of n=8 naïve mice was used for comparison. For this test, radiant heat was used to induce withdrawal responses, with stimulation directed at the hindpaw (Plantar Test Apparatus, IITC Life Science). The right and left paws were each tested 3-5 times, with at least 5 minutes between replicate measurements, and the closest two values from each set (i.e. per paw) were averaged to obtain withdrawal latencies. These were then averaged over both paws to compute overall thermal sensitivity.

### Thermal Preference Test

The Thermal Preference test was performed one day following the Hargreaves test. Mice were placed in an individual rectangular enclosure whose flooring was a thermal plate with temperature gradient controlled by Peltier heating and cooling elements across the length of the enclosure (constructed by William N. Goolsby). Tests applied a gradient range of 20° -40° C to the thermal platform, centered around the mouse thermoneutral preference point of ∼30° C.^37^ Because the platform temperature followed a linear gradient, preferred temperature could be calculated using the animal’s body position (midline [rostro-caudal and left-right] location). Mice were allowed to explore for 30 minutes to 1 hour and their chosen resting spot after this time period was taken to indicate their individual thermal preference temperature.

### Statistical Analysis

All quantitative measurements are reported as mean ± SEM. Analysis of variance (one-way ANOVA, with follow-up multiple comparisons tests) was used for respiratory measures and activity, a paired t-test to assess pre-post von Frey score changes, and an unpaired t-test to assess thermal gradient preferences relative to a separate cohort of naïve mice. To quantify the relationship between sensory tests and physiological variables (respiration and movement), correlation analyses were undertaken at timepoints corresponding to observed changes in SCI mice, with linear regression performed to assess statistical significance. To avoid statistical over-testing, correlations were performed based on the *a priori* hypothesis that the earliest changes after SCI would provide the greatest opportunity for revealing predictive relationships relevant to clinical translation and preventative interventions. Comparisons were made between changes in respiration and activity at acute (1 dpo) and subacute (8 dpo) timepoints and development of pain-related outcomes at 4 weeks post-SCI. In the case of significant results, further analysis was performed at 2 and 3 dpo to provide added clarification on the timeframe of effects. Statistics were performed using GraphPad Prism software, Version 10.2.1 for Windows (GraphPad Software, Inc.; San Diego, CA), with significance set at p<.05 and two-tailed tests.

## Results

This study investigated if early changes in respiration and activity could be used to predict the development of neuropathic pain after SCI.

### RR, RRVAR, and activity levels

As shown in **Figure 2A**, RR did not significantly change from the baseline within the first 8 days after operation (one-way RM ANOVA; F(1.945, 23.34)=0.1257, p>.05). In contrast, RRVAR (**Figure 2B**) changed after injury (one-way RM ANOVA; F(2.370, 28.44)=8.927, p<.001), with significantly decreased values on the first two days after injury compared to baseline (p<0.05, Dunnett’s multiple comparisons tests). RRVAR recovered by the 8^th^ day post injury. Time spent moving (**Figure 2C**) also changed in the days after injury (mixed effects model due to one missing data point; p<0.0001), with post-hoc tests revealing a significant decrease on each of the first three days after injury (p<.0001). Like RRVAR, time spent moving recovered by the 8^th^ day post injury. While there was a borderline significant correlation (p = 0.05) between decreases in RRVAR and percent time active from baseline to 1 dpo, these variables were not correlated within individual mice when averaged across 1-3 dpo (**Figure 2D)**.

**Figure 2:**
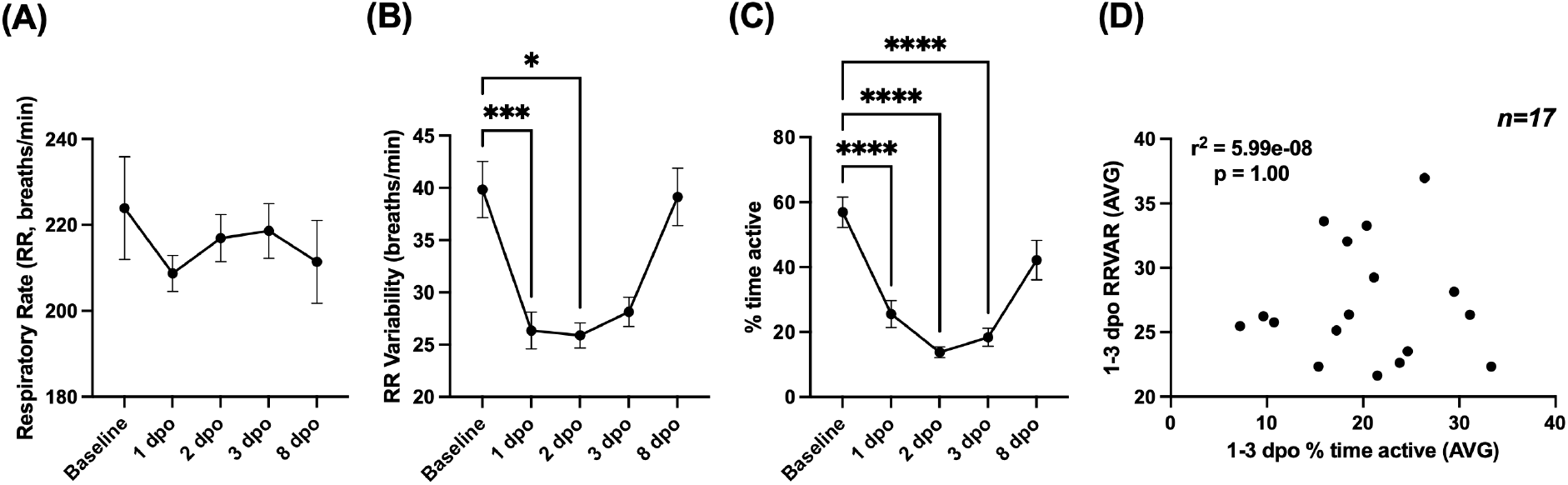
Comparing respiratory rate (RR), RR variability (RRVAR), and movement changes in the early stages after SCI. These measures were recorded from one day pre-injury (baseline) to 8 days post-injury in spinal cord injured (SCI) mice (n=17, except 2 dpo [n=13]). (**A)** RR did not change significantly throughout the experiment. (**B)** RRVAR significantly decreased on the first and second days post operation compared to baseline and recovered by 8 dpo. (**C)** Time spent moving significantly decreased on each of the first 3 dpo and recovered by 8 dpo. For A-C, * p<.05, ^**^ p<.005, and ^****^ p<.0001; post-hoc tests after 1-factor RM ANOVA. (**D)** Average percent time active and RRVAR from 1-3 dpo were uncorrelated across all mice. dpo, days post operation.

### Mechanical and thermal pain assessments

The von Frey test was conducted at baseline and 4 weeks post-SCI, and the Hargreaves at 4 weeks post-SCI, to assess changes in threshold for paw withdrawal reflexes in response to mechanical and heat stimuli, respectively. As seen in previously studies,^19,31,34^ the threshold filament force needed to induce a withdrawal response in the von Frey test decreased by four weeks after SCI, consistent with mechanical allodynia (**Figure 3A**; paired t-test; t(9)=7.356, p<0.0001).

**Figure 3:**
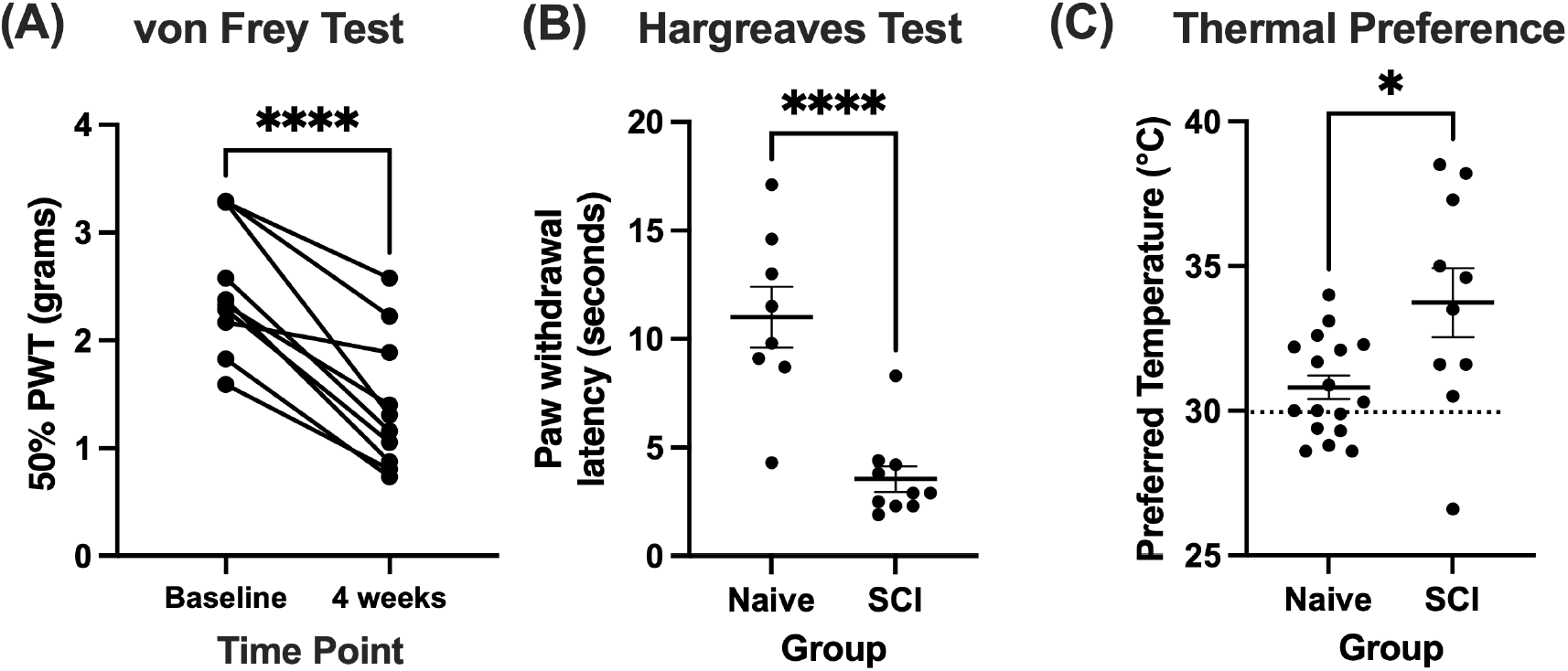
Von Frey test scores and thermal preferences. *N=10 SCI* (**A**) von Frey scores for all mice tested at baseline (1 day pre-injury) and 4-weeks post-injury. The lines between data points indicate changes within individual mice. There was a statistically significant change in von Frey scores 4 weeks post-injury (**** p<.0001). PWT, paw withdrawal threshold (**B**) The Hargreaves Test for heat allodynia was run at 4 weeks post-injury in SCI mice. SCI mice withdrew their hindpaws significantly earlier than a control population of naïve mice (*N=8*; unpaired t-test, **** p<.0001). (**C**) The thermal preference test across a temperature gradient of 20-40° C, undertaken at 4 weeks post-injury. The average thermal preference after SCI was significantly higher than the thermoneutral temperature in naïve mice (*N=17*; unpaired t-test, * p=.01).

In the Hargreaves test (**Figure 3B**), mice had a mean latency of 3.55±0.59s to remove their hindpaw from the thermal stimulus, which was significantly lower than the withdrawal latency of 11.01±1.40s in a control cohort of n=8 naïve mice (t(16)=5.296, p<.0001), consistent with heat allodynia.

In the Thermal Preference test, conducted at 4 weeks post-SCI (**Figure 3C**), SCI mice preferred a temperature of 33.7±1.2° C. This value was higher than the reported thermoneutrality point of 30° C^37^ and significantly higher than the preferred temperature of 30.8 ± 0.4° C observed in a control cohort of n=17 naïve mice (unpaired t-test; t(25)=2.785, p=0.01), suggesting thermal dysregulation.

### Correlations between early SCI changes and later pain-related outcomes

To understand the relationship between early measures of RRVAR and movement with the subsequent development of dysfunction, correlation analyses were performed. We assessed whether acute and subacute changes after injury were associated with changes in measured mechanical and thermal sensitivity. A comprehensive table of correlations is included in **Table 1**. Larger decreases in RRVAR from baseline to 1 day after SCI corresponded to less sensitivity to stimulation in the von Frey test four weeks after injury (greater withdrawal thresholds, p=0.01). Follow up analysis revealed that this effect was also significant from baseline to 3 dpo (p=0.024) with the same trend at 2 dpo (p=0.06). We also investigated the relationship between early changes in RRVAR after SCI and future behavioral outcomes (**Figure 4**). The decrease in RRVAR at 1 dpo (compared to its recovery at 8 dpo) was positively correlated with increases in temperature preferences at 4 weeks post-injury (**Figure 4A**), i.e. early reductions in RRVAR that recovered by 8 dpo predicted a preference for hotter temperatures. Follow-up analyses revealed that this effect was also significant at 2 dpo (p=0.009) and 3 dpo (p=0.01) vs. the 8 dpo timepoint.

**Table 1:**
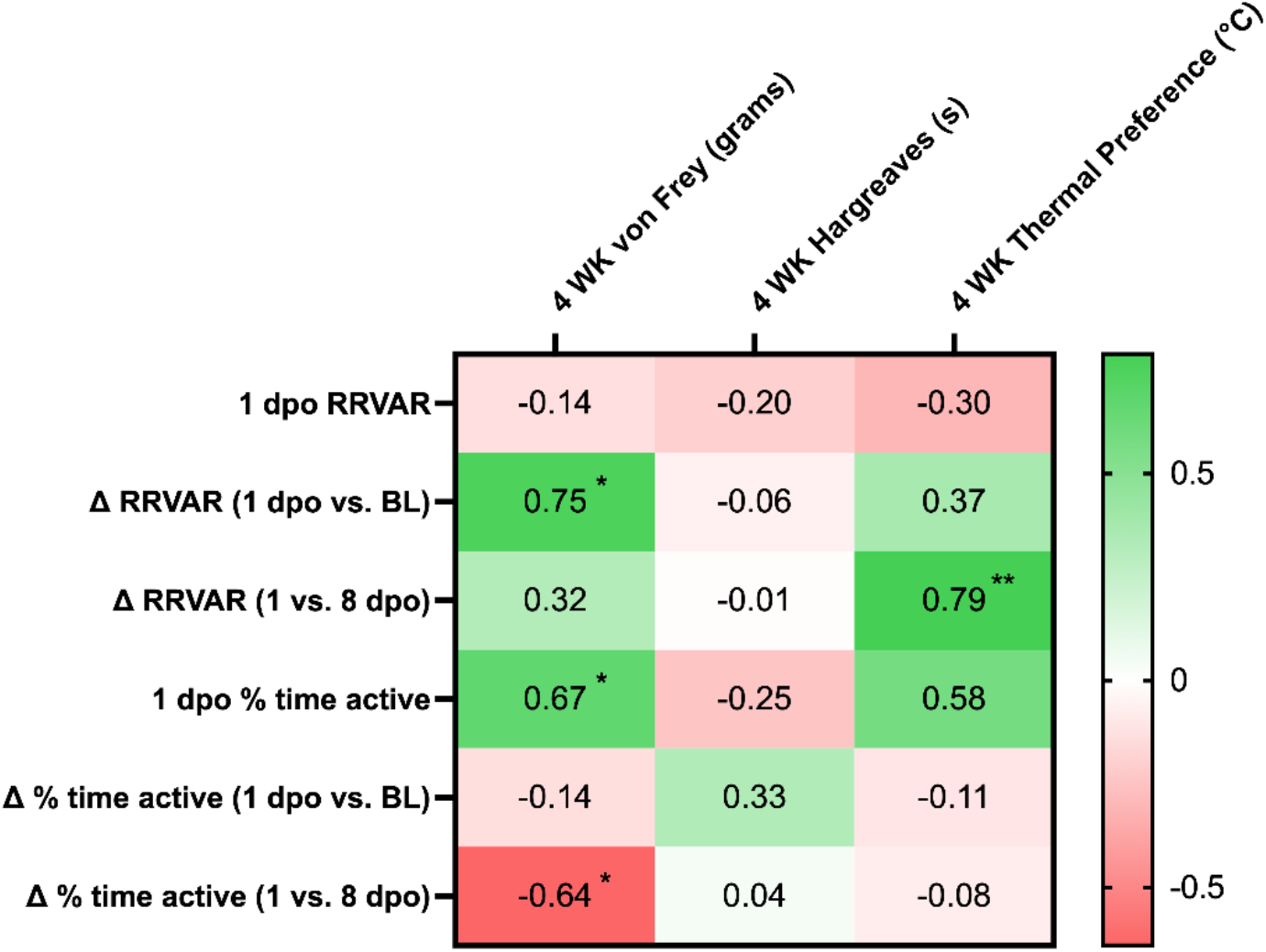
Correlations between independent (RRVAR and activity levels) and dependent (pain-related behaviors) variables. Pearson correlation coefficient (r) values were obtained to assess whether acute and subacute changes after injury were associated with pain-related outcomes. To avoid statistical over-testing (i.e. multiple comparisons problems), correlations were performed based on the a priori hypothesis that the earliest changes after SCI would provide the greatest opportunity for revealing predictive relationships relevant to outcomes at the chronic stage. Comparisons were made between changes in RRVAR and activity at acute (1 dpo) and subacute (8 dpo) timepoints and development of pain-and temperature-related outcomes at 4 weeks post-SCI. Greener colors indicate greater positive correlations and redder colors negative correlations. Significance values and interpretation are provided in the manuscript text. Note that **Figure 2** shows statistical comparisons to baseline, since our first goal was to assess behavioral change after SCI. In contrast, correlations assessed 1 dpo changes compared to baseline as well as to 8 dpo (**Figure 4**) to understand how the evolution of early deficits predicted chronic dysfunction. *Δ RRVAR indicates the absolute value of the difference between 1 dpo vs. BL or 1 vs. 8dpo*. ^*\**^ *p<*.*05*, ^*\*\**^ *p<*.*01*

**Figure 4:**
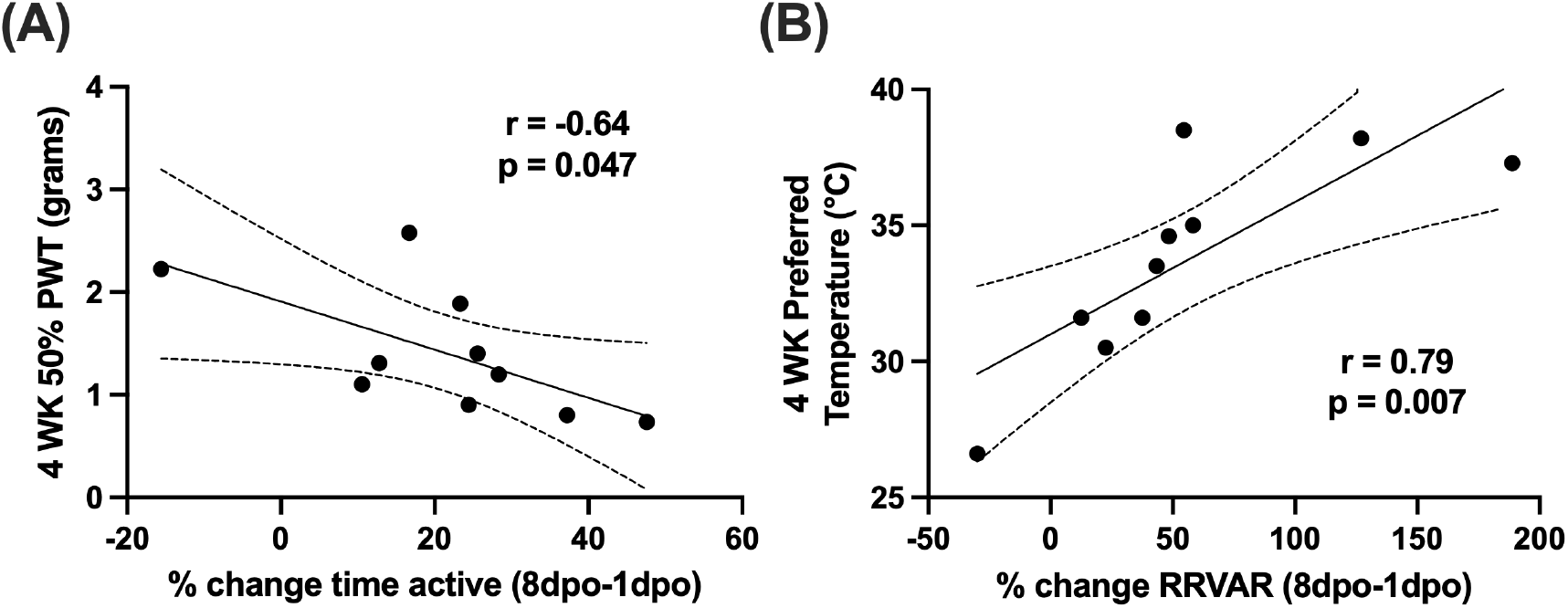
Correlating early recovery of movement and RR variability (RRVAR) with pain outcomes. **(A)** Decreased % time active at 1 compared to 8 dpo was negatively correlated with von Frey scores (i.e., emergent mechano-sensitivity) at 4 weeks post-injury. **(B)** Decreases in RRVAR at 1 compared to 8 dpo were significantly associated with higher preferred floor temperatures at 4 weeks post-injury. *N = 10*; dpo, days post injury; BL, baseline; RR, respiratory rate; RR VAR, respiratory rate variability; PWT, paw withdrawal threshold

Exploring the relationship between movement and future hypersensitivity, recovery of activity (change in % time active from 1 to 8 dpo) was negatively correlated with von Frey scores at 4 weeks post-injury (**Figure 4B**), i.e. animals with greater recovery showed more mechanical hypersensitivity. Follow up analyses revealed that there was a significant correlation between reduction in % time active at 1 dpo and 4-week von Frey score (p=0.036) but this correlation was lost at 2 and 3 dpo.

## Discussion

This study captured RR, RRVAR, and percentage of time spent moving before and at several time points after a contusion SCI. Consistent with our previous reports in mice,^31^ RR did not change 24 hours after SCI. In contrast, we previously found positive correlations between early changes in RR and mechanical hypersensitivity in rats.^19^ This may be explainable by differences in severity between the hemisection and contusion SCI models used. Here, RRVAR decreased steeply from 1-3 days after SCI. The early changes in RRVAR mirrored early decreases in movement and eventual recovery by 8 dpo. While there was a borderline significant correlation between the magnitude of decrease in these two variables from baseline to 1 dpo – potentially reflecting injury severity – absolute RRVAR and movement levels were uncorrelated over the first three days post-injury. Furthermore, deficits in RRVAR and movement over the first week after SCI differentially predicted the emergence of mechano-sensitivity and higher temperature thermal place preference. In contrast, none of the early changes after SCI were predictive of subsequent emergence of heat allodynia.

After chronic SCI, animals demonstrated heat allodynia and increased preference for higher temperatures in the thermal gradient. The latter effect may indicate a compromise in thermoregulation or the emergence of cold allodynia.^3,38-40^ The normal mouse thermoneutral point of ∼30ºC is also the highest temperature to trigger peripheral cold receptors in Aδ- and C-fibers,^41^ so observed mouse preferences for temperatures above 30ºC may support the emergence of cold allodynia. While T10 contusion does not present with the same degree of thermoregulatory dysfunction seen at higher-level SCIs,^42,43^ the elevated thermal preference of ∼34ºC is consistent with a compromise in thermoregulation associated with impaired activation of below-SCI level skin sympathetic vasoconstrictors.^44,45^ In contrast, heat allodynia results from activation of heat receptors at temperatures higher than ∼40ºC,^46,47^ well above observed SCI mouse preferences. Future SCI studies would benefit from more precise tests on heat and cold allodynia^48^ in relation to measures of thermal place preference.

Correlation analyses revealed that larger decreases in RRVAR one day after SCI compared to baseline predicted decreased mechanical hypersensitivity at 4 weeks. This early change did not predict future changes in thermal preference or heat pain. In contrast, differences in RRVAR between 1 and 8 dpo correlated with a preference for higher temperatures 4 weeks after SCI, implying that greater RRVAR recovery predicted greater thermal dysfunction. Though both RRVAR and thermoregulatory status are controlled by autonomic circuits, it is unclear how their neural circuitry would interact.^45^ More complete assessment of time-dependent changes in autonomic variables after SCI is warranted. Overall, the magnitude of SCI-induced early RRVAR decrease vs RRVAR time-dependent magnitude changes in recovery differentially predicted mechanical hypersensitivity and thermal preference, respectively.

As with RRVAR, early post-SCI changes in movement did not predict emergence of heat pain. In contrast, we found a significant correlation between reductions in activity at 1 dpo and the magnitude of mechanical allodynia. Mice with the greatest activity decreases at 1 versus 8 dpo were the most likely to develop mechano-sensitivity (see **Table 1**). Together, these results suggest that tracking the trajectories of activity and RRVAR over the first eight days following an SCI provides complementary information predicting development of chronic dysfunction. While the negative relationship between RRVAR decreases at 1 dpo and mechanical allodynia was an unexpected finding, it could suggest that immediate autonomic responses after SCI may reflect other physiological changes (e.g. increased parasympathetic activity^49,50^) that are protective against later development of tactile allodynia. In contrast, decreased activity levels at 1 dpo predicted development of hypersensitivity in the von Frey test four weeks post-injury, supporting broad locomotor impairment as an early predictor of risk for mechanical pain.

SCI presents with relatively well documented early deficits in locomotion^51^ and inflammation,^52^ and our understanding of acute-to-chronic sequelae leading to neuropathic pain has gradually improved.^53^ However, identifying early predictors of emerging pain phenotypes remains an unmet need.^5^ Several studies have suggested potential biomarkers for SCI chronic neuropathic pain.^40,54-56^ Peripheral hypersensitivity two weeks to one month after injury can predict development of central pain in humans,^40,57^ a time period which may approximately correspond to the first 24 hours in mice.^58^ Our laboratory previously found positive correlations between spontaneous primary afferent activity after SCI and RRVAR at later time points.^18^ However, to our knowledge this is the first study to demonstrate respiratory and motor deficits in the first 24 hours after injury that predict behavioral deficits at a chronic time point.

In conclusion, we found pronounced deficits in RRVAR and activity for 1-3 days following a lower thoracic SCI in mice, with accompanying mechanical allodynia and thermal dysregulation at a chronic time point. Activity levels 24 hours after SCI and the evolution of RRVAR in the first week post-injury differentially predicted which animals went on to develop chronic changes in behavior. Future studies are needed to better understand the effect that these and other physiological measures have on SCI chronic pain symptoms, for instance by using a machine learning approach on raw biosensor data. It will also be important to test for cold allodynia^59-61^ and/or autonomic nervous system dysfunction by incorporating pharmacological, behavioral, and physiological approaches.^62,63^ Overall, this study provides insights on predicting behavioral dysfunction after SCI and suggests future research avenues for clarifying the complex causal mechanisms underlying neuropathic pain.

## Acknowledgements

The authors would like to thank William N. Goolsby for his help with device design and construction, and for technical support.

## Authors’ Contributions

Conception or design of the work: S.H., D.J.N., and A.C. Acquisition of data for the work: A.C. and D.J.N. Analysis and interpretation of data for the work: D.J.N., A.C., and S.H. Drafting the work: A.C., D.J.N., and S.H. Reviewing the work critically for important intellectual content: A.C., D.J.N., and S.H. Final approval of the version to be published: A.C., D.J.N., and S.H. Agreement to be accountable for all aspects of the work in ensuring that questions related to the accuracy or integrity of any part of the work are appropriately investigated and resolved: A.C., D.J.N., and S.H.

## References

1. Baastrup C, Finnerup NB. Pharmacological management of neuropathic pain following spinal cord injury. CNS Drugs 2008;22(6):455–75

2. Siddall P. Spinal cord injury pain: a retrospective. In: Spinal Cord Injury Pain. (Sang CN, Hulsebosch CE. eds.) Academic Press: Cambridge, MA; 2022; pp. 25–43.

3. Shiao R, Lee-Kubli CA. Neuropathic Pain After Spinal Cord Injury: Challenges and Research Perspectives. Neurotherapeutics 2018;15(3):635–653, doi:10.1007/s13311-018-0633-4

4. Colloca L, Ludman T, Bouhassira D, et al. Neuropathic pain. Nat Rev Dis Primers 2017;3(17002, doi:10.1038/nrdp.2017.2

5. Widerstrom-Noga E. Neuropathic Pain and Spinal Cord Injury: Phenotypes and Pharmacological Management. Drugs 2017;77(9):967–984, doi:10.1007/s40265-017-0747-8

6. Kinnman E, Levine JD. Sensory and sympathetic contributions to nerve injury-induced sensory abnormalities in the rat. Neuroscience 1995;64(3):751–67, doi:10.1016/0306-4522(94)00435-8

7. Kinnman E, Levine JD. Involvement of the sympathetic postganglionic neuron in capsaicin-induced secondary hyperalgesia in the rat. Neuroscience 1995;65(1):283–91, doi:10.1016/0306-4522(94)00474-j

8. Jänig W, Levine JD, Michaelis M. Chapter 10. Interactions of sympathetic and primary afferent neurons following nerve injury and tissue trauma. In: Progress in Brain Research. (Takao Kumazawa LK, Kazue M. eds.) Elsevier: 1996; pp. 161–184.

9. Ramer MS, Thompson SWN, McMahon SB. Causes and consequences of sympathetic basket formation in dorsal root ganglia. Pain 1999;82, Supplement 1(0):S111–S120, doi:http://dx.doi.org/10.1016/S0304-3959(99)00144-X

10. Zhang J-M, Li H, Munir MA. Decreasing sympathetic sprouting in pathologic sensory ganglia: a new mechanism for treating neuropathic pain using lidocaine. Pain 2004;109(1– 2):143-149, doi:http://dx.doi.org/10.1016/j.pain.2004.01.033

11. Loggia ML, Juneau M, Bushnell MC. Autonomic responses to heat pain: Heart rate, skin conductance, and their relation to verbal ratings and stimulus intensity. Pain 2011;152(3):592–8, doi:10.1016/j.pain.2010.11.032

12. Culman J, Ritter S, Ohlendorf C, et al. A new formalin test allowing simultaneous evaluation of cardiovascular and nociceptive responses. Can J Physiol Pharmacol 1997;75(10-11):1203-11

13. Santuzzi CH, Neto Hde A, Pires JG, et al. High-frequency transcutaneous electrical nerve stimulation reduces pain and cardio-respiratory parameters in an animal model of acute pain: participation of peripheral serotonin. Physiother Theory Pract 2013;29(8):630–8, doi:10.3109/09593985.2013.774451

14. Wolf S, Hardy JD. Studies on Pain. Observations on Pain Due to Local Cooling and on Factors Involved in the “Cold Pressor” Effect. J Clin Invest 1941;20(5):521–33, doi:10.1172/JCI101245

15. Hodgetts TJ, Kenward G, Vlachonikolis IG, et al. The identification of risk factors for cardiac arrest and formulation of activation criteria to alert a medical emergency team. Resuscitation 2002;54(2):125–131, doi:Pii S0300-9572(02)00100-4 Doi 10.1016/S0300-9572(02)00100-4

16. Fieselmann JF, Hendryx MS, Helms CM, et al. Respiratory rate predicts cardiopulmonary arrest for internal medicine inpatients. Journal of general internal medicine 1993;8(7):354–60

17. Subbe CP, Davies RG, Williams E, et al. Effect of introducing the Modified Early Warning score on clinical outcomes, cardio-pulmonary arrests and intensive care utilisation in acute medical admissions. Anaesthesia 2003;58(8):797–802, doi:10.1046/j.1365-2044.2003.03258.x

18. Idlett-Ali S, Kloefkorn H, Goolsby W, et al. Relating Spinal Injury-Induced Neuropathic Pain and Spontaneous Afferent Activity to Sleep and Respiratory Dysfunction. J Neurotrauma 2023;40(23-24):2654–2666, doi:10.1089/neu.2022.0305

19. Noble DJ, Martin KK, Parvin S, et al. Spontaneous and Stimulus-Evoked Respiratory Rate Elevation Corresponds to Development of Allodynia in Spinal Cord-Injured Rats. J Neurotrauma 2019, doi:10.1089/neu.2018.5936

20. Noble DJ, Hochman S. Hypothesis: Pulmonary Afferent Activity Patterns During Slow, Deep Breathing Contribute to the Neural Induction of Physiological Relaxation. Front Physiol 2019;10(1176, doi:10.3389/fphys.2019.01176

21. Jensen MP, Kuehn CM, Amtmann D, et al. Symptom burden in persons with spinal cord injury. Arch Phys Med Rehabil 2007;88(5):638–45, doi:10.1016/j.apmr.2007.02.002

22. Bryson M, Kloefkorn H, Idlett-Ali S, et al. Emergent epileptiform activity in spinal sensory circuits drives ectopic bursting in afferent axons and sensory dysfunction after cord injury. Pain 2025;166(2):e27–e35, doi:10.1097/j.pain.0000000000003364

23. Pfyffer D, Vallotton K, Curt A, et al. Tissue bridges predict neuropathic pain emergence after spinal cord injury. J Neurol Neurosurg Psychiatry 2020;91(10):1111–1117, doi:10.1136/jnnp-2020-323150

24. Thornton WA, Marzloff G, Ryder S, et al. The presence or absence of midsagittal tissue bridges and walking: a retrospective cohort study in spinal cord injury. Spinal Cord 2023;61(8):436–440, doi:10.1038/s41393-023-00890-6

25. Xu ML, Wu XB, Liang Y, et al. A Silver Lining of Neuropathic Pain: Predicting Favorable Functional Outcome in Spinal Cord Injury. J Pain Res 2023;16(2619-2632, doi:10.2147/JPR.S414638

26. Metz GA, Curt A, van de Meent H, et al. Validation of the weight-drop contusion model in rats: a comparative study of human spinal cord injury. J Neurotrauma 2000;17(1):1–17, doi:10.1089/neu.2000.17.1

27. Young W. Spinal cord contusion models. Prog Brain Res 2002;137(231-55, doi:10.1016/s0079-6123(02)37019-5

28. Murakami T, Kanchiku T, Suzuki H, et al. Anti-interleukin-6 receptor antibody reduces neuropathic pain following spinal cord injury in mice. Exp Ther Med 2013;6(5):1194–1198, doi:10.3892/etm.2013.1296

29. Basso DM, Fisher LC, Anderson AJ, et al. Basso Mouse Scale for locomotion detects differences in recovery after spinal cord injury in five common mouse strains. J Neurotrauma 2006;23(5):635–59, doi:10.1089/neu.2006.23.635

30. Jakeman LB, Chen Y, Lucin KM, et al. Mice lacking L1 cell adhesion molecule have deficits in locomotion and exhibit enhanced corticospinal tract sprouting following mild contusion injury to the spinal cord. Eur J Neurosci 2006;23(8):1997–2011, doi:10.1111/j.1460-9568.2006.04721.x

31. Noble DJ, Dongmo R, Parvin S, et al. C-low threshold mechanoreceptor activation becomes sufficient to trigger affective pain in spinal cord-injured mice in association with increased respiratory rates. Front Integr Neurosci 2022;16(1081172, doi:10.3389/fnint.2022.1081172

32. Noble DJ, Goolsby WN, Garraway SM, et al. Slow Breathing Can Be Operantly Conditioned in the Rat and May Reduce Sensitivity to Experimental Stressors. Front Physiol 2017;8(854, doi:10.3389/fphys.2017.00854

33. Noble DJ, MacDowell CJ, McKinnon ML, et al. Use of electric field sensors for recording respiration, heart rate, and stereotyped motor behaviors in the rodent home cage. J Neurosci Methods 2017;277(88-100, doi:10.1016/j.jneumeth.2016.12.007

34. Garraway SM, Woller SA, Huie JR, et al. Peripheral noxious stimulation reduces withdrawal threshold to mechanical stimuli after spinal cord injury: role of tumor necrosis factor alpha and apoptosis. Pain 2014;155(11):2344–59, doi:10.1016/j.pain.2014.08.034

35. Chaplan SR, Bach FW, Pogrel JW, et al. Quantitative assessment of tactile allodynia in the rat paw. Journal of neuroscience methods 1994;53(1):55–63

36. Hargreaves K, Dubner R, Brown F, et al. A new and sensitive method for measuring thermal nociception in cutaneous hyperalgesia. Pain 1988;32(1):77–88

37. Fischer AW, Cannon B, Nedergaard J. Optimal housing temperatures for mice to mimic the thermal environment of humans: An experimental study. Mol Metab 2018;7(161-170, doi:10.1016/j.molmet.2017.10.009

38. Rossi HL, Vierck CJ, Jr., Caudle RM, et al. Characterization of cold sensitivity and thermal preference using an operant orofacial assay. Mol Pain 2006;2(37, doi:10.1186/1744-8069-2-37

39. Yoon YW, Dong H, Arends JJ, et al. Mechanical and cold allodynia in a rat spinal cord contusion model. Somatosens Mot Res 2004;21(1):25–31, doi:10.1080/0899022042000201272

40. Finnerup NB, Norrbrink C, Trok K, et al. Phenotypes and predictors of pain following traumatic spinal cord injury: a prospective study. J Pain 2014;15(1):40–8, doi:10.1016/j.jpain.2013.09.008

41. McKemy DD. TRPM8: The Cold and Menthol Receptor. In: TRP Ion Channel Function in Sensory Transduction and Cellular Signaling Cascades. (Liedtke WB, Heller S. eds.) Boca Raton (FL); 2007.

42. Alilain WJ, Horn KP, Hu H, et al. Functional regeneration of respiratory pathways after spinal cord injury. Nature 2011;475(7355):196–200, doi:10.1038/nature10199

43. Sharma H, Alilain WJ, Sadhu A, et al. Treatments to restore respiratory function after spinal cord injury and their implications for regeneration, plasticity and adaptation. Experimental neurology 2012;235(1):18–25, doi:10.1016/j.expneurol.2011.12.018

44. Price MJ, Trbovich M. Thermoregulation following spinal cord injury. Handb Clin Neurol 2018;157(799-820, doi:10.1016/B978-0-444-64074-1.00050-1

45. Jänig W. The integrative action of the autonomic nervous system : neurobiology of homeostasis. Cambridge University Press: Cambridge, United Kingdom; New York, NY; 2022.

46. Dubin AE, Patapoutian A. Nociceptors: the sensors of the pain pathway. J Clin Invest 2010;120(11):3760–72, doi:10.1172/JCI42843

47. Zimmermann K, Hein A, Hager U, et al. Phenotyping sensory nerve endings in vitro in the mouse. Nat Protoc 2009;4(2):174–96, doi:10.1038/nprot.2008.223

48. Gao T, Hao J, Wiesenfeld-Hallin Z, et al. Activation of TRPM8 cold receptor triggers allodynia-like behavior in spinally injured rats. Scand J Pain 2013;4(1):33–37, doi:10.1016/j.sjpain.2012.09.007

49. Henke AM, Billington ZJ, Gater DR, Jr. Autonomic Dysfunction and Management after Spinal Cord Injury: A Narrative Review. J Pers Med 2022;12(7), doi:10.3390/jpm12071110

50. Teasell RW, Arnold JM, Krassioukov A, et al. Cardiovascular consequences of loss of supraspinal control of the sympathetic nervous system after spinal cord injury. Arch Phys Med Rehabil 2000;81(4):506–16, doi:10.1053/mr.2000.3848

51. Duan WR, Huang Q, Chen ZY, et al. Comparisons of motor and sensory abnormalities after lumbar and thoracic contusion spinal cord injury in male rats. Neuroscience letters 2019;708(doi:ARTN 134358 10.1016/j.neulet.2019.134358

52. Rice T, Larsen J, Rivest S, et al. Characterization of the early neuroinflammation after spinal cord injury in mice. J Neuropath Exp Neur 2007;66(3):184–195, doi:DOI 10.1097/01.jnen.0000248552.07338.7f

53. Gaudet AD, Ayala MT, Schleicher WE, et al. Exploring acute-to-chronic neuropathic pain in rats after contusion spinal cord injury. Experimental neurology 2017;295(46-54, doi:10.1016/j.expneurol.2017.05.011

54. Gruener H, Zeilig G, Gaidukov E, et al. Biomarkers for predicting central neuropathic pain occurrence and severity after spinal cord injury: results of a long-term longitudinal study. Pain 2020;161(3):545–556, doi:10.1097/j.pain.0000000000001740

55. Vuckovic A, Gallardo VJF, Jarjees M, et al. Prediction of central neuropathic pain in spinal cord injury based on EEG classifier. Clin Neurophysiol 2018;129(8):1605–1617, doi:10.1016/j.clinph.2018.04.750

56. Levitan Y, Zeilig G, Bondi M, et al. Predicting the Risk for Central Pain Using the Sensory Components of the International Standards for Neurological Classification of Spinal Cord Injury. J Neurotrauma 2015;32(21):1684–92, doi:10.1089/neu.2015.3947

57. Zeilig G, Enosh S, Rubin-Asher D, et al. The nature and course of sensory changes following spinal cord injury: predictive properties and implications on the mechanism of central pain. Brain 2012;135(Pt 2):418–30, doi:10.1093/brain/awr270

58. Dutta S, Sengupta P. Men and mice: Relating their ages. Life Sci 2016;152(244-8, doi:10.1016/j.lfs.2015.10.025

59. Ruan Y, Gu L, Yan J, et al. An effective and concise device for detecting cold allodynia in mice. Sci Rep 2018;8(1):14002, doi:10.1038/s41598-018-31741-7

60. Deuis JR, Dvorakova LS, Vetter I. Methods Used to Evaluate Pain Behaviors in Rodents. Front Mol Neurosci 2017;10(284, doi:10.3389/fnmol.2017.00284

61. MacDonald DI, Wood JN, Emery EC. Molecular mechanisms of cold pain. Neurobiol Pain 2020;7(100044, doi:10.1016/j.ynpai.2020.100044

62. Rabchevsky AG, Kitzman PH. Latest approaches for the treatment of spasticity and autonomic dysreflexia in chronic spinal cord injury. Neurotherapeutics 2011;8(2):274–82, doi:10.1007/s13311-011-0025-5

63. Noble DJ, MacDowell CJ, McKinnon ML, et al. Use of electric field sensors for recording respiration, heart rate, and stereotyped motor behaviors in the rodent home cage. Journal of neuroscience methods 2017;277(88–100)

